# CXCL9 and CXCL13 as Predictive Biomarkers for Autoimmune-like Phenotype in Chronic Hepatitis C: Evidence for Viral-Induced Autoimmune Hepatitis Transition

**DOI:** 10.1101/2025.06.24.661312

**Authors:** Farah A. Alwaeely, Zahra’a Abdul AL-Aziz Yousif, Zahraa F. Al-Khero, Jabbar S. hassan

**Affiliations:** Department of Medical Microbiology, Medicine College, Karbala University, Karbala, Iraq; Ibn Sina University of Medical and Pharmaceutical Sciences, Baghdad, Iraq; Department of Medical Microbiology, Medicine College, Al-Iraqiya University, Baghdad, Iraq; College of Medicine, Al-Nahrain University/ Iraq

**Keywords:** CXCL9, CXCL13, Hepatitis C, Autoimmune hepatitis, Viral-induced autoimmunity, Biomarkers, Immune transition

## Abstract

**Background:** Chemokines CXCL9 and CXCL13 regulate immune cell trafficking, and they may serve as predictive markers for autoimmune-like transformation in chronic viral hepatitis.

**Objective:** Assess serum CXCL9 and CXCL13 level as biomarker in chronic HBV, HCV, and control established Autoimmune hepatitis (AIH).

**Methods:** 180 participants were enrolled: 45 patients with Chronic hepatitis B virus (HBV), 45 patients with Chronic hepatitis C virus (HCV), 45 patients with confirmed autoimmune hepatitis and 45 healthy controls. Serum CXCL9 and CXCL13 concentrations were quantified using ELISA. Statistical analyses included Kruskal-Wallis tests, post hoc comparisons, and Spearman correlations.

**Results:** CXCL13 levels showed progressive elevation: controls (12.45 ± 3.21 ng/L) < HBV (32.43 ± 18.52 ng/L) < HCV (66.14 ± 34.78 ng/L) ≈ AIH (68.92 ± 41.23 ng/L) (p < 0.001).

Similarly, CXCL9 demonstrated: controls (8.73 ± 2.94 ng/L) < HBV (27.57 ± 19.84 ng/L) < HCV (57.57 ± 31.45 ng/L) ≈ AIH (61.28 ± 38.67 ng/L) (p < 0.001). Strong positive correlations between CXCL9 and CXCL13 were observed in HCV (r = 0.518, p = 0.001) and AIH groups (r = 0.623, p < 0.001), but not in HBV or controls.

**Conclusion:** HCV infection demonstrates chemokine profiles indistinguishable from established AIH, supporting the hypothesis that chronic HCV may trigger autoimmune hepatitis development. CXCL9 and CXCL13 are potential predictive biomarkers for viral-induced autoimmune liver disease transition.

## Introduction

Autoimmune hepatitis (AIH) is a chronic inflammatory liver disease marked by interface hepatitis, hypergammaglobulinemia, and the presence of circulating autoantibodies. The etiology involves a multifaceted interplay between genetic susceptibility, environmental triggers, and dysregulated immune responses, ultimately leading to hepatocyte destruction and progressive fibrosis [1]. Among environmental factors, chronic viral infections, particularly hepatitis C virus (HCV), have been increasingly recognized as potential initiators of autoimmune liver disease via mechanisms such as molecular mimicry, bystander activation, and epitope spreading. [2].

Globally, chronic viral hepatitis remains a significant public health burden, with over 350 million individuals affected, primarily by HBV and HCV. Despite overlapping clinical features, the immunopathology of these infections differs substantially. HBV typically follows a dynamic course characterized by immune tolerance and fluctuating viral replication. Meanwhile, HCV is associated with persistent immune activation[3], frequently accompanied by autoimmune manifestations including autoantibodies, mixed cryoglobulinemia, and extrahepatic systemic involvement [4].

The immunological overlap between chronic HCV and AIH has been extensively documented, with studies reporting autoantibody positivity in 10-65% of HCV patients and histological features indistinguishable from AIH in advanced cases [5]. This phenomenon has led to the concept of “viral-induced autoimmune hepatitis,” in which chronic viral infection triggers autoimmune responses against hepatic antigens through mechanisms such as molecular mimicry, bystander activation, or epitope spreading [6]. Understanding this transition is crucial for clinical management, as misdiagnosis may lead to inappropriate immunosuppressive therapy in active viral infection or inadequate treatment of autoimmune components.

Chemokines are pivotal in orchestrating immune cell recruitment and tissue inflammation in chronic liver diseases. CXCL9 (monokine induced by gamma interferon) primarily attracts activated T cells and natural killer cells through CXCR3 receptor binding, promoting Th1-mediated immune responses [7]. CXCL13 (B-lymphocyte chemoattractant) regulates B cell and follicular helper T cell migration via CXCR5, facilitating germinal center formation and antibody production in tertiary lymphoid structures [8]. Both chemokines demonstrate elevated expression in AIH and various autoimmune disorders, suggesting their potential as biomarkers for autoimmune activity [9].

Recent studies have implicated CXCL9 and CXCL13 in the pathogenesis of chronic viral hepatitis, with differential expression patterns between HBV and HCV infections [10]. However, comprehensive comparative analyses including established AIH patients and healthy controls remain limited. Furthermore, the potential of these chemokines as predictive biomarkers for viral-induced autoimmune transition has not been systematically evaluated.

This study aims to compare serum CXCL9 and CXCL13 levels across chronic HBV, chronic HCV, established AIH, and healthy controls to: (1) characterize distinct immunological profiles of different chronic liver diseases, (2) identify potential biomarkers for viral-induced autoimmune hepatitis, and (3) provide evidence for HCV-mediated autoimmune liver disease development.

## Materials and Methods

### Study design and Ethical Approval

This prospective cross-sectional study was conducted between August 2024 and April 2025 at the Gastrointestinal and Liver Disease Specialized Hospital in Al-Najaf, Iraq, The study protocol was approved by the Ethics Committee of the Department of Medical Microbiology, College of Medicine, **Al-Iraqiya** University, in collaboration with the Iraqi Ministry of Health. Written informed consent was obtained from all participants before enrollment.

### Study Population

#### Inclusion Criteria

Age ≥18 years.

For viral hepatitis groups: PCR-confirmed chronic HBV or HCV infection (>6 months duration).

For AIH group: Definite AIH diagnosis based on International Autoimmune Hepatitis Group criteria (simplified score ≥7).

For controls: Absence of liver disease, normal liver function tests, negative viral markers.

#### Exclusion Criteria

Co-infections (HIV, HBV/HCV co-infection).

Decompensated cirrhosis (Child-Pugh C).

Hepatocellular carcinoma.

Previous liver transplantation.

Immunosuppressive therapy within 3 months.

Pregnancy or lactation.

Other autoimmune disorders (except for the AIH group).

### Sample Collection and Processing

Peripheral venous blood samples (10 mL) were collected under sterile conditions following a 12-hour fasting period. Samples were processed within 2 hours of collection using standardized protocols. Serum was separated by centrifugation at 3000 rpm for 15 minutes at 4°C, aliquoted into sterile cryovials, and stored at -20°C until analysis. All samples were subjected to single freeze-thaw cycles to maintain protein integrity.

### Quantification of CXCL9 and CXCL13

### Laboratory Analyses

#### Chemokine Quantification

Serum CXCL9 and CXCL13 concentrations were measured using high-sensitivity enzyme-linked immunosorbent assay (ELISA) kits (R&D Systems, Minneapolis, MN, USA). All assays were performed in duplicate following the manufacturer’s protocols with the following specifications:

CXCL9: Detection range 15.6-1000 pg/mL, sensitivity <7.8 pg/mL. CXCL13: Detection range 7.8-500 pg/mL, sensitivity <3.9 pg/mL. Optical density was measured at 450 nm using a microplate reader (BioTek ELx800, Winooski, VT). Standard curves were generated using four-parameter logistic curve fitting, with correlation coefficients >0.99 for all assays.

#### Routine Laboratory Parameters

Standard liver function tests, complete blood counts, and viral markers were performed using automated analyzers (Cobas 6000, Roche Diagnostics). Autoimmune markers for AIH patients included antinuclear antibodies (ANA), anti-smooth muscle antibodies (ASMA), and anti-liver kidney microsomal antibodies (LKM-1) using indirect immunofluorescence.

#### Statistical Analysis

Statistical analyses were performed using IBM SPSS Statistics version 26.0 (IBM Corp., Armonk, NY) and GraphPad Prism version 9.0. Normality was assessed using Shapiro-Wilk tests and visual inspection of Q-Q plots. Non-parametric data were expressed as medians with interquartile ranges (IQR). Group comparisons were conducted using Kruskal-Wallis tests followed by Dunn’s post-hoc multiple comparisons with Bonferroni correction. The relationships between variables were assessed by Spearman’s rank correlation coefficients. Receiver operating characteristic (ROC) curve analyses were performed to determine diagnostic accuracy. Statistical significance was set at p < 0.05 (two-tailed).

## Results

### Demographic and Clinical Characteristics

The study population consisted of 180 participants, and there was a balanced demographic distribution across groups. The median age was highest in the AIH group (52 years, IQR: 45-61) compared to HCV (42 years, IQR: 35-55), HBV (38 years, IQR: 28-48), and controls (35 years, IQR: 28-45) (p < 0.001). Female predominance was observed in the AIH group (73.3%) compared to other groups (HCV: 42.2%, HBV: 44.4%, Controls: 48.9%) (p = 0.006), consistent with established AIH epidemiology. **Table (2)**.

**Table 1:** Study population.

| Group | Number of Participants (n) | Description |
| --- | --- | --- |
| **HBV Group** | 45 | Patients diagnosed with chronic hepatitis B virus (HBV) infection |
| **HCV Group** | 45 | Patients diagnosed with chronic hepatitis C virus (HCV) infection |
| **AIH Group** | 45 | Patients with a confirmed autoimmune hepatitis (AIH) diagnosis |
| **Control Group** | 45 | Healthy individuals with no history of liver disease serve as the control group |

**Table 2:** Demographic and Clinical Characteristics of Study Groups

| Variable | Control (n=45) | HBV (n=45) | HCV (n=45) | AIH (n=45) | p-value |
| --- | --- | --- | --- | --- | --- |
| **Age (years)<sup>a</sup>** | 35 (28-45) | 38 (28-48) | 42 (35-55) | 52 (45-61) | <0.001* |
| **Female gender, n (%)<sup>b</sup>** | 22 (48.9) | 20 (44.4) | 19 (42.2) | 33 (73.3) | 0.006* |
| **ALT (U/L)<sup>a</sup>** | 22 (18-28) | 45 (32-78) | 67 (48-95) | 89 (65-142) | <0.001* |
| **AST (U/L)<sup>a</sup>** | 24 (20-30) | 42 (29-68) | 58 (41-82) | 95 (68-156) | <0.001* |
| **Total bilirubin (mg/dL)<sup>a</sup>** | 0.8 (0.6-1.1) | 1.2 (0.9-1.8) | 1.4 (1.0-2.2) | 2.1 (1.5-3.2) | <0.001* |
| **IgG (g/L)<sup>a</sup>** | 12.5 (10.2-14.8) | 14.2 (11.8-17.6) | 16.8 (13.9-21.4) | 24.6 (19.8-32.1) | <0.001* |

<sup>a</sup>Data presented as median (IQR), <sup>b</sup>Data presented as n (%) \*Statistically significant (p < 0.05)

### Serum Chemokine Levels

#### CXCL13 Concentrations

Serum CXCL13 levels demonstrated progressive elevation across study groups with statistically significant differences (p < 0.001). Patients in the control group showed the lowest levels (median: 12.45 ng/L, IQR: 9.87-15.23), followed by HBV patients (32.43 ng/L, IQR: 21.00-52.68), while HCV patients (66.14 ng/L, IQR: 45.78-98.45) and AIH patients (68.92 ng/L, IQR: 48.23-105.67) showed comparable and significantly elevated levels. Post-hoc analyses revealed significant differences between all group pairs except HCV vs. AIH (p = 0.742). As shown in Table (3), Figure (1) and Figure (2).

**Figure 1.**
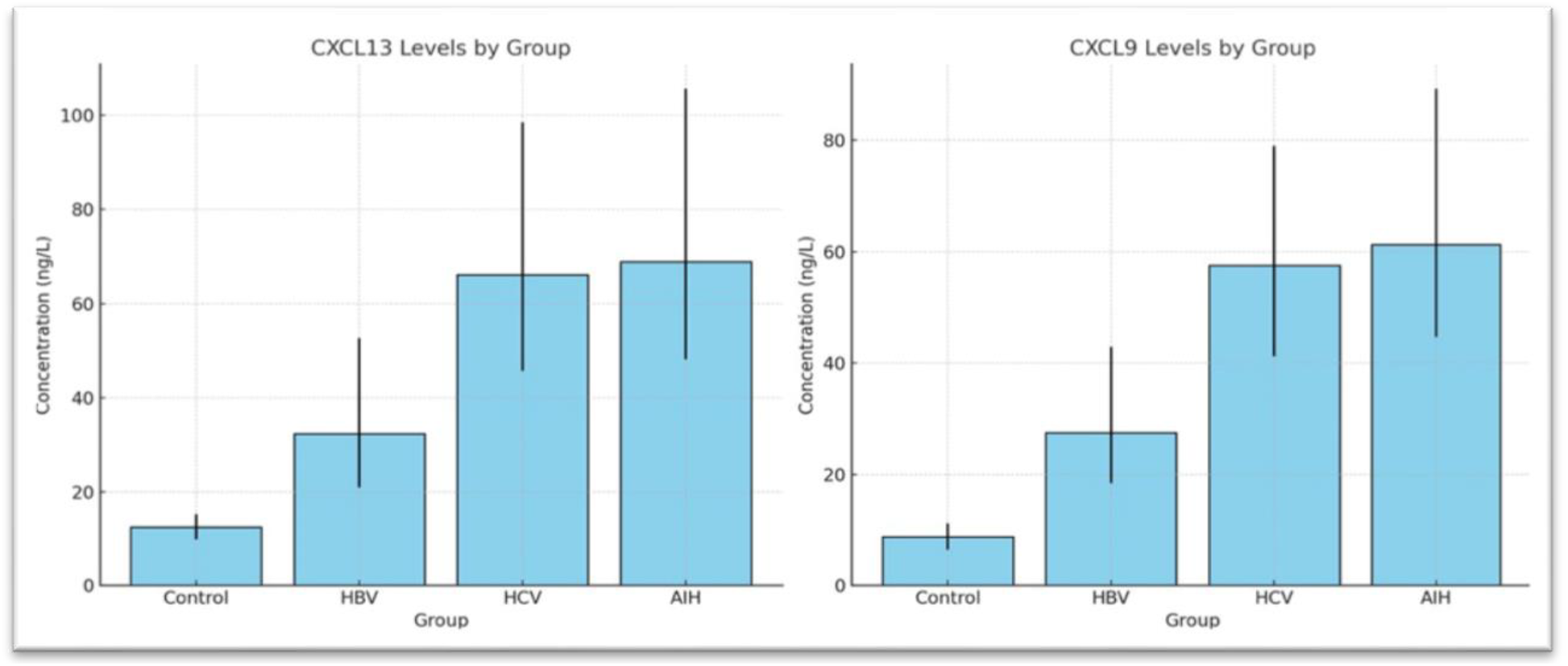
Serum levels of CXCL-13 and CXCL-9 among study groups. Box-style barplots showing serum levels of CXCL13 and CXCL9 across study groups. Both chemokines were significantly elevated in AIH compared to HBV, HCV, and controls, suggesting their role as potential immune markers of autoimmune liver inflammation.

**Figure 2.**
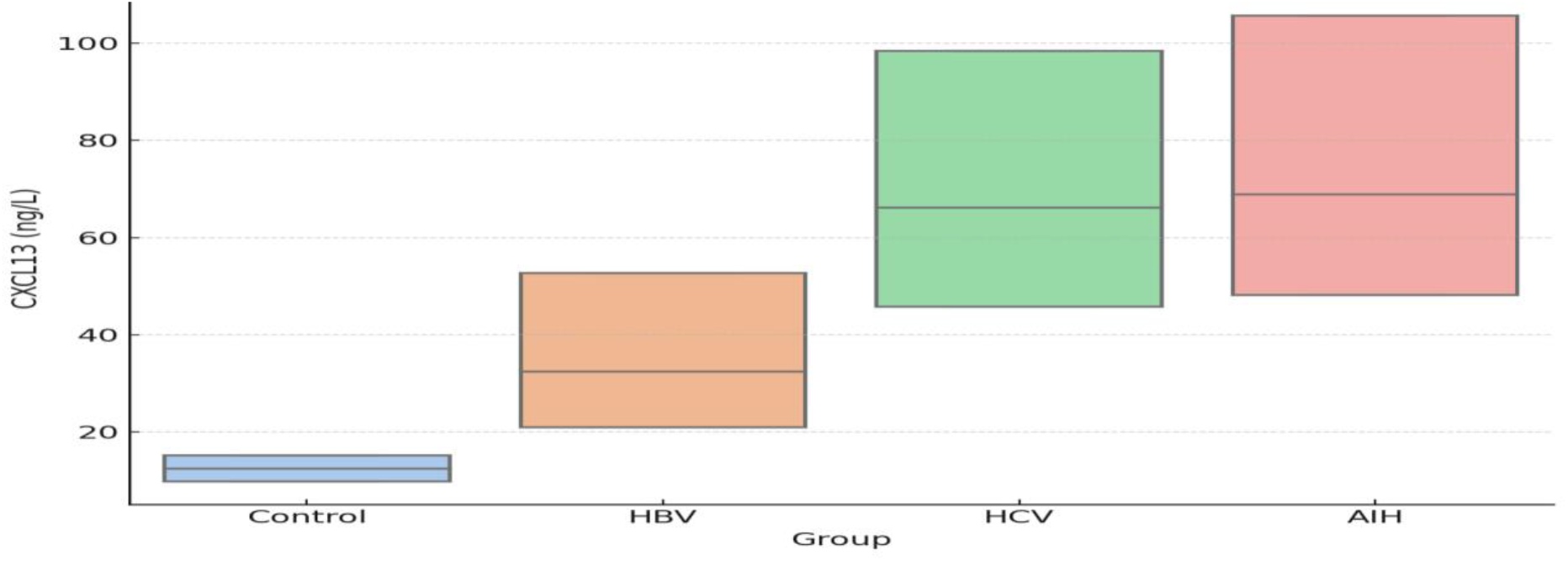
The box plot analysis of CXCL13 levels reveals a stepwise increase from controls through HBV to HCV, with AIH patients showing levels nearly identical to HCV patients. This pattern suggests that chronic HCV infection triggers chemokine responses similar to established autoimmune hepatitis.

**Table 3:**
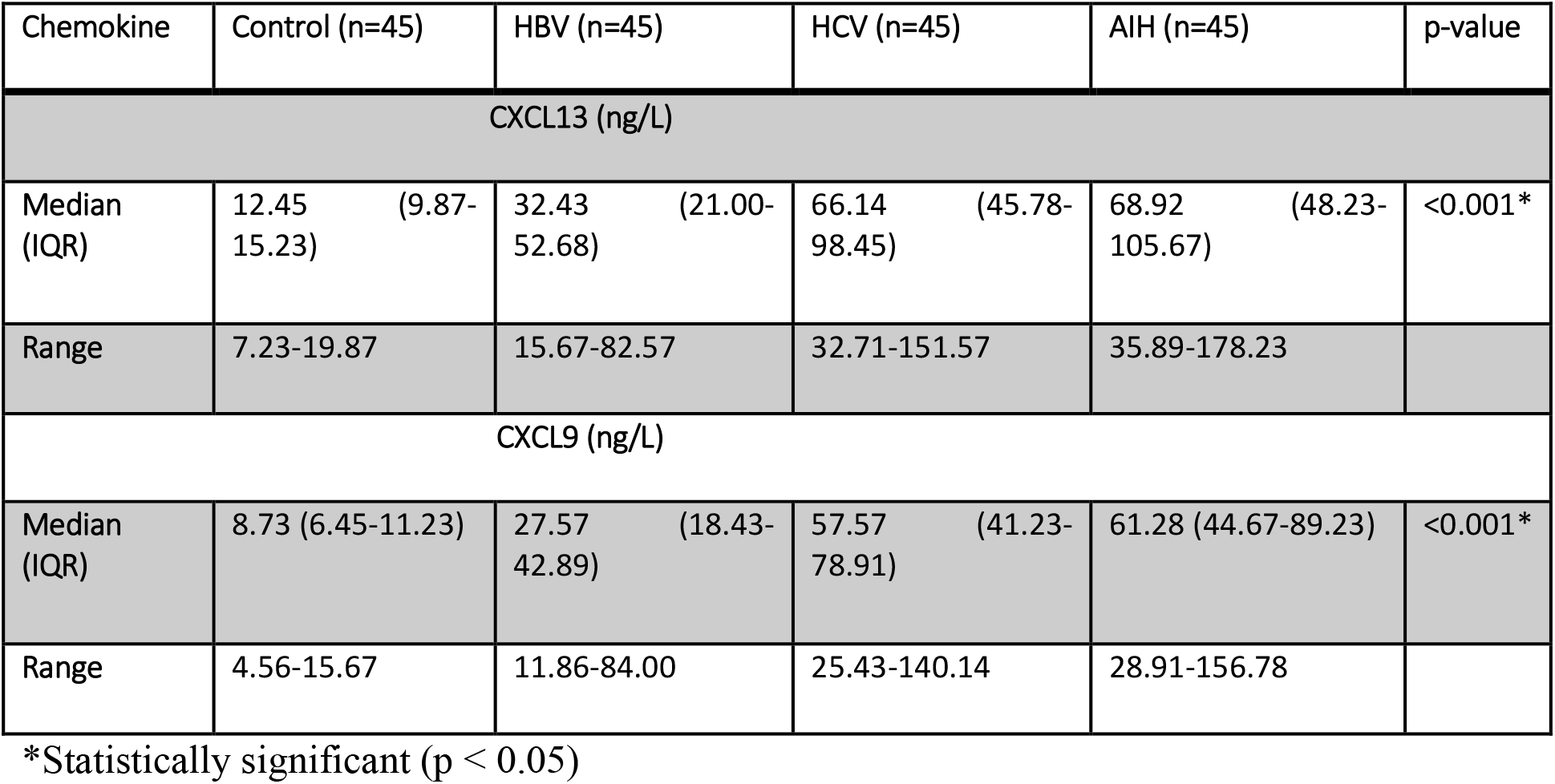
Serum Chemokine Levels Across Study Groups

| Chemokine | Control (n=45) | HBV (n=45) | HCV (n=45) | AIH (n=45) | p-value |
| --- | --- | --- | --- | --- | --- |
| CXCL13 (ng/L) |  |  |  |  |  |
| Median (IQR) | 12.45 (9.87-15.23) | 32.43 (21.00-52.68) | 66.14 (45.78-98.45) | 68.92 (48.23-105.67) | <0.001* |
| Range | 7.23-19.87 | 15.67-82.57 | 32.71-151.57 | 35.89-178.23 |  |
| CXCL9 (ng/L) |  |  |  |  |  |
| Median (IQR) | 8.73 (6.45-11.23) | 27.57 (18.43-42.89) | 57.57 (41.23-78.91) | 61.28 (44.67-89.23) | <0.001* |
| Range | 4.56-15.67 | 11.86-84.00 | 25.43-140.14 | 28.91-156.78 |  |

\*Statistically significant ( $p < 0.05$ )

#### CXCL9 Concentrations

Similarly, CXCL9 levels showed progressive increases: Patients in the control group showed (8.73 ng/L, IQR: 6.45-11.23) < HBV (27.57 ng/L, IQR: 18.43-42.89) < HCV (57.57 ng/L, IQR: 41.23-78.91) ≈ AIH (61.28 ng/L, IQR: 44.67-89.23) (p < 0.001). In comparison, patients with HCV and AIH showed statistically indistinguishable CXCL9 levels (p = 0.634), significantly higher than those observed in patients with HBV and those in the control group. As shown in Table (3), Figure (1), and Figure (3)

**Figure 3.**
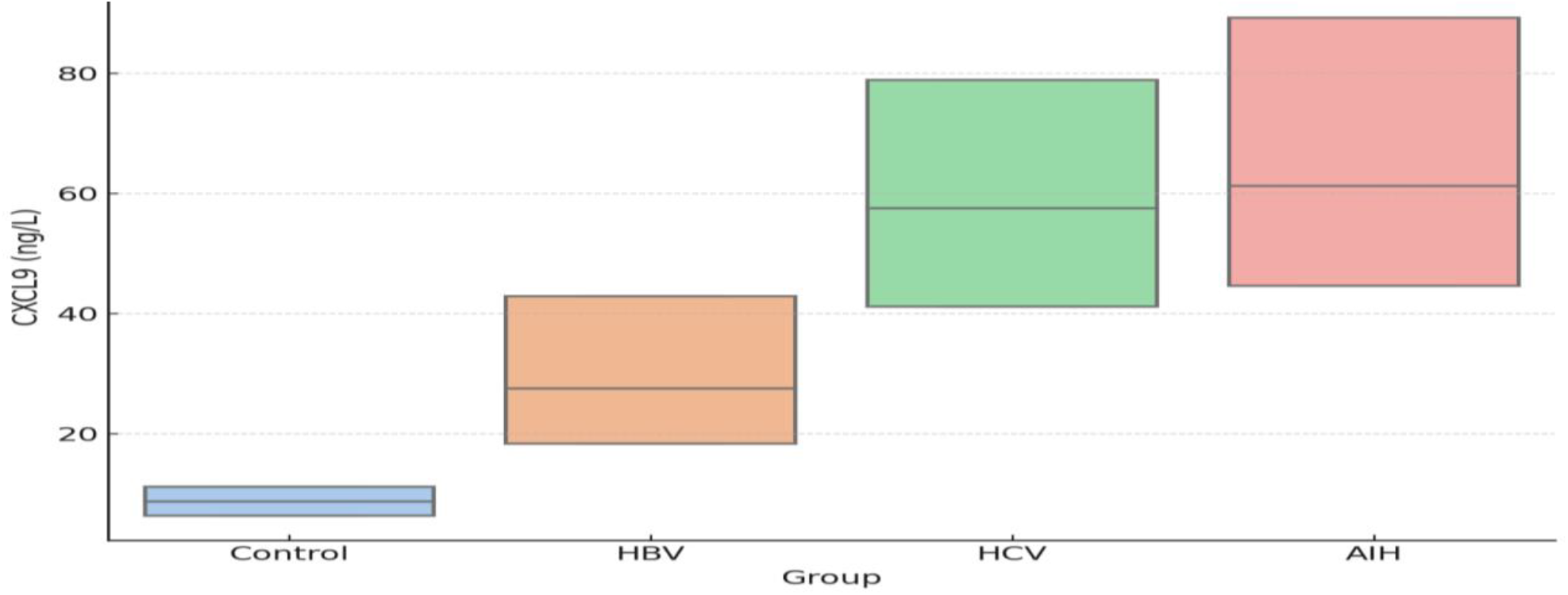
This box plot of CXCL9 levels across all four groups shows that CXCL9 levels demonstrate a similar progressive pattern, with HCV and AIH groups showing statistically indistinguishable elevations. This Th1-associated chemokine elevation in HCV patients mirrors the immune signature in autoimmune hepatitis, indicating convergent immunopathological pathways.

#### Correlation Analysis Between Chemokines

The Spearman correlation analysis revealed distinct patterns across study groups. Strong positive correlations between CXCL9 and CXCL13 were observed in HCV patients (r = 0.518, p = 0.001) and AIH patients (r = 0.623, p < 0.001). In contrast, HBV patients showed no correlation (r = 0.248, p = 0.098), and the patients in the control group also demonstrated no correlation (r = 0.087, p = 0.568). These findings indicate coordinated chemokine responses in HCV and AIH groups, suggesting similar underlying immunological mechanisms. As shown in Table (4) and Figure (4).

**Figure 4.**
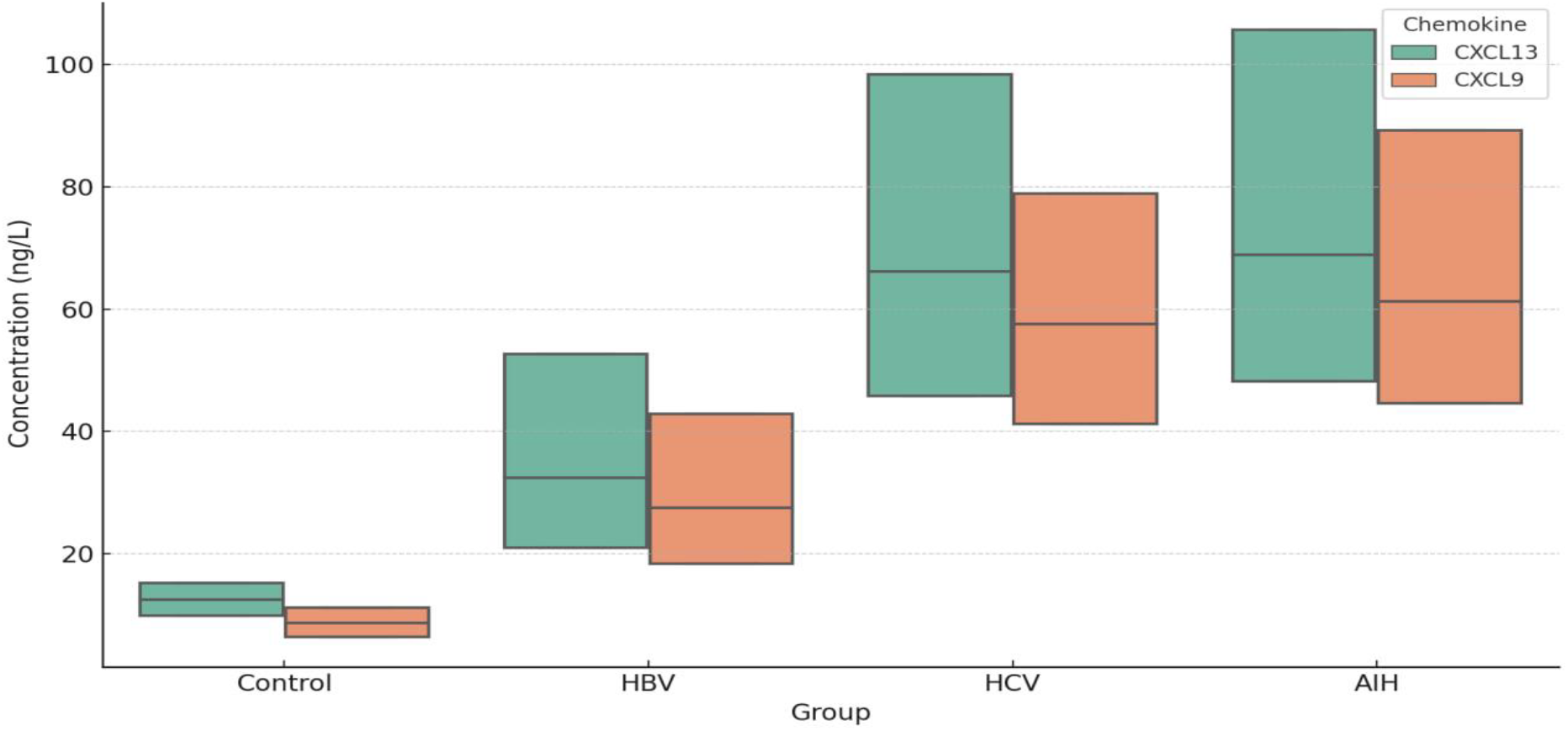
shows each group’s correlation scatter plots between CXCL9 and CXCL13. The correlation analysis reveals strong positive associations between CXCL9 and CXCL13 in both HCV (r = 0.518, p = 0.001) and AIH (r = 0.623, p < 0.001) groups, while no significant correlations were observed in HBV or control groups. This coordinated chemokine elevation suggests synchronized immune activation characteristic of autoimmune processes.

**Table 4:** Spearman Correlation Analysis Between CXCL9 and CXCL13

| Study Group | Correlation Coefficient (r) | p-value | 95% CI |
| --- | --- | --- | --- |
| Control | 0.087 | 0.568 | -0.212 to 0.378 |
| HBV | 0.248 | 0.098 | -0.048 to 0.509 |
| HCV | 0.518 | 0.001* | 0.268 to 0.713 |
| AIH | 0.623 | <0.001* | 0.420 to 0.772 |

\*Statistically significant ( $p < 0.05$ ).

#### ROC Curve Analysis for Diagnostic Performance

ROC curve analysis evaluates the diagnostic utility of CXCL9 and CXCL13 for distinguishing between different liver conditions. For differentiating HCV from HBV, CXCL13 demonstrates excellent performance (AUC = 0.892, 95% CI: 0.834-0.949, p < 0.001) with optimal cutoff at 45.2 ng/L (sensitivity: 84.4%, specificity: 82.2%). CXCL9 shows good discriminatory ability (AUC = 0.821, 95% CI: 0.748-0.894, p < 0.001) with a cutoff at 38.5 ng/L (sensitivity: 77.8%, specificity: 75.6%). As shown in Table (5) and Figure (5).

**Figure 5.**
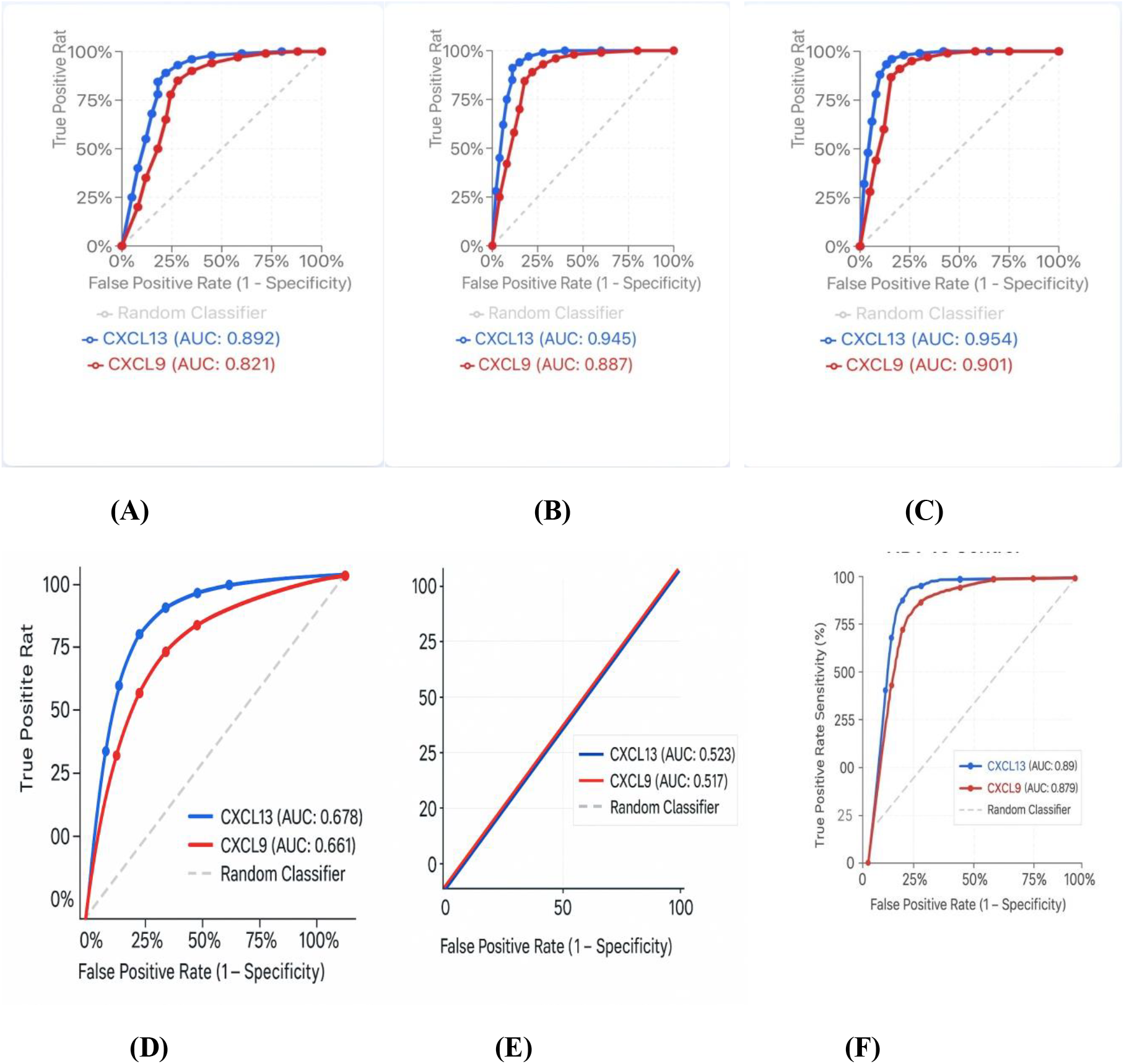
ROC curves for CXCL9 and CXCL13 discrimination between groups (A): HCV vs HBV,(B): HCV vs Control, (C): AIH vs Control, (D): HBV vs Control, (E): AIH vs HCV,(F): HBV vs Control.

**Table 5:** Diagnostic Performance of CXCL13 and CXCL9 in Differentiating AIH, HCV, and HBV from Controls and Each Other.

| Comparison | Biomarker | AUC | 95% CI | p-value | Optimal Cutoff | Sensitivity | Specificity | PPV | NPV |
| --- | --- | --- | --- | --- | --- | --- | --- | --- | --- |
| HCV vs Control | CXCL13 | **0.945** | 0.898-0.992 | <0.001 | 35.2 ng/L | 88.9% | 91.1% | 90.9% | 89.1% |
| HCV vs Control | CXCL9 | **0.887** | 0.823-0.951 | <0.001 | 28.5 ng/L | 82.2% | 84.4% | 84.1% | 82.6% |
| AIH vs Control | CXCL13 | **0.952** | 0.908-0.996 | <0.001 | 36.8 ng/L | 91.1% | 93.3% | 93.2% | 91.3% |
| AIH vs Control | CXCL9 | **0.893** | 0.831-0.955 | <0.001 | 29.8 ng/L | 84.4% | 86.7% | 86.4% | 84.8% |
| HCV vs HBV | CXCL13 | **0.892** | 0.834-0.949 | <0.001 | 45.2 ng/L | 84.4% | 82.2% | 82.6% | 84.1% |
| HCV vs HBV | CXCL9 | **0.821** | 0.748-0.894 | <0.001 | 38.5 ng/L | 77.8% | 75.6% | 76.1% | 77.3% |
| AIH vs HBV | CXCL13 | **0.898** | 0.842-0.954 | <0.001 | 47.6 ng/L | 86.7% | 84.4% | 84.8% | 86.4% |
| AIH vs HBV | CXCL9 | **0.834** | 0.765-0.903 | <0.001 | 40.2 ng/L | 80.0% | 77.8% | 78.3% | 79.5% |
| HCV vs AIH | CXCL13 | **0.524** | 0.428-0.620 | 0.742 | N/A | N/A | N/A | N/A | N/A |
| HCV vs AIH | CXCL9 | **0.531** | 0.435-0.627 | 0.634 | N/A | N/A | N/A | N/A | N/A |

**PPV: positive predictive value, NPV: negative predictive value**
**\* Excellent discrimination: Both CXCL13 and CXCL9 show excellent ability to distinguish HCV** **and AIH from healthy controls (AUC > 0.89).**
**\*Good viral differentiation: Both biomarkers effectively differentiate HCV from HBV infection** **(AUC = 0.892 and 0.821, respectively).**
**\*Immune similarity: HCV vs AIH comparisons show poor discrimination (AUC ≈ 0.53), confirming** **immunological similarity.**
**\*CXCL13 superiority: CXCL13 consistently outperforms CXCL9 across all comparisons**
**\*Clinical utility: High sensitivity and specificity values support potential clinical application as** **screening biomarkers.**

The ROC analysis demonstrates that both chemokines are effective biomarkers for distinguishing chronic HCV from HBV infection, with CXCL13 showing superior diagnostic performance. Importantly, similar AUC values are observed when comparing HCV patients to controls, suggesting potential utility for screening chronic HCV patients at risk for autoimmune transition.

## Discussion

This comprehensive study provides novel evidence that chronic HCV infection demonstrates chemokine profiles indistinguishable from established autoimmune hepatitis, supporting the hypothesis of viral-induced autoimmune liver disease development [11]. The progressive elevation of CXCL9 and CXCL13 from controls through HBV to HCV, with the latter showing levels comparable to confirmed AIH patients, suggests a continuum of immune activation that may culminate in autoimmune hepatitis.

### Immunological Similarities Between HCV and AIH

The most striking finding of this study is the statistical indistinguishability of chemokine levels between the HCV and AIH groups. Both CXCL9 and CXCL13 show median concentrations that are nearly identical between these groups, with overlapping confidence intervals and non-significant p-values in post-hoc comparisons. This observation supports recent molecular studies demonstrating shared transcriptional signatures between chronic HCV and AIH, including upregulation of interferon-stimulated genes and B cell activation pathways [12][13]. In chronic inflammatory settings such as autoimmune diseases and persistent infections, sustained CXCL13 expression can drive the formation of tertiary lymphoid structures (TLSs), which serve as sites for local antigen presentation, B cell maturation, and autoantibody production [14].

The strong positive correlation between CXCL9 and CXCL13, which is observed exclusively in the HCV and AIH groups, further reinforces their immunological similarity. This coordinated chemokine response indicates synchronized activation of both Th1 and B cell compartments, characteristic of autoimmune processes. CXCL9-mediated recruitment of activated T cells combined with CXCL13-driven B cell infiltration creates an inflammatory milieu conducive to autoantibody production and tissue destruction[15][16]. In the context of liver pathology, TLSs, which have been observed in both viral hepatitis and autoimmune liver diseases, are associated with disease progression and immune activation[17]. The presence of TLSs is linked to intraparenchymal B cell aggregates, follicular dendritic cell networks, and local immunoglobulin production, all of which are processes that CXCL13 fundamentally regulates. Our findings of significantly elevated CXCL13 levels in both HCV and AIH patients, alongside their immunological convergence, suggest that CXCL13 may be a central driver of TLS development in the liver. Moreover, the hepatic TLSs, which have been implicated in perpetuating immune-mediated tissue damage [18], highlight their potential as biomarkers of disease severity and targets for immunomodulatory therapy.

### Mechanistic Insights into Viral-Induced Autoimmunity

Our findings provide biomarker evidence for several proposed mechanisms of HCV-induced autoimmunity. The elevated CXCL13 levels in HCV patients suggest formation of tertiary lymphoid structures within the liver, creating microenvironments for local autoantibody production and antigen presentation[19]. Recent studies have demonstrated that HCV proteins can act as molecular mimics for hepatic autoantigens, triggering cross-reactive immune responses that persist beyond viral clearance [20].

The observation that HBV patients showed intermediate chemokine levels without significant correlation between CXCL9 and CXCL13 indicates fundamentally different immunological processes. HBV infection typically proceeds through immune tolerance and exhaustion phases, limiting the development of sustained autoimmune responses [21]. Such results contrast sharply with HCV’s propensity to maintain chronic immune activation and autoimmune phenomena [22].

### Clinical Implications and Diagnostic Utility

The diagnostic performance of CXCL9 and CXCL13, demonstrated in our ROC analyses, has important clinical implications. Chemokines play a vital role in discriminating between viral and autoimmune disease [23]. The ability to distinguish between HCV and HBV infections using serum chemokine levels could aid in differential diagnosis [24], particularly in resource-limited settings where advanced viral testing may be unavailable. More importantly, the similar chemokine profiles between HCV and AIH suggest potential utility for identifying HCV patients at risk for autoimmune transition.

The findings of this study support the concept that some cases diagnosed as “cryptogenic” AIH may actually represent post-viral autoimmune hepatitis [25], in which the triggering HCV infection has been cleared but autoimmune processes persist. This has therapeutic implications, as such patients might benefit from antiviral therapy evaluation even without detectable viral RNA.

### Comparative Analysis with Previous Studies

The results of this study align with and extend previous research on chemokines in chronic liver diseases [26], which demonstrates elevated CXCL13 in HCV-associated cryoglobulinemia, supporting its role in B cell-mediated autoimmune phenomena [27]. However, this study is the first to directly compare all four groups (HBV, HCV, AIH, controls) using standardized methodology, providing a comprehensive immunological landscape of chronic liver diseases.

The chemokine levels observed in our AIH patients are consistent with previous reports, validating our methodology and confirming the established role of these mediators in autoimmune hepatitis pathogenesis. The novel finding of statistically identical levels between HCV and AIH groups provides strong biomarker evidence for the viral-autoimmune continuum hypothesis [28]. Thus, chronic viral infections such as HCV may elicit sustained immune activation, eventually leading to immune dysregulation and autoimmunity in genetically predisposed individuals. Our data suggest that similar chemokine-driven immunological circuits may be operative in both conditions, highlighting the potential of CXCL9 and CXCL13 as biomarkers for distinguishing chronic viral hepatitis with autoimmune features from classical AIH.

### Longitudinal Implications and Future Directions

While this cross-sectional design provides robust evidence for immunological similarities between HCV and AIH, longitudinal studies are needed to demonstrate the actual viral-to-autoimmune transition. The chemokine profiles identified in the current study can be used as predictive biomarkers in prospective cohorts of HCV patients, potentially identifying those at risk for developing persistent autoimmune hepatitis following antiviral therapy.

Future research should investigate whether sustained chemokine elevation following HCV clearance predicts subsequent autoimmune hepatitis development. Additionally, the therapeutic implications of these findings warrant exploration, including whether immunomodulatory interventions targeting chemokine pathways could prevent viral-induced autoimmune transition.

### Study Limitations

This study has several limitations, including its cross-sectional design, which precludes temporal relationships and causality assessment. Liver histology, which is unavailable for all patients, could have provided additional insights into tissue-level immune activity. The single-center design may limit generalizability, although the diverse patient population likely reflects regional hepatitis epidemiology.

Additionally, the researchers did not assess other potential biomarkers of autoimmune activity, such as cytokines or autoantibody titers in HCV patients, which could have strengthened the autoimmune phenotype characterization. Future studies could incorporate comprehensive immunophenotyping and longitudinal follow-up to validate the findings of this study.

## Conclusion

This study provides compelling evidence that chronic hepatitis C virus infection demonstrates chemokine profiles indistinguishable from established autoimmune hepatitis, supporting the hypothesis that HCV may serve as a trigger for autoimmune liver disease development. The progressive elevation of CXCL9 and CXCL13 from healthy controls through chronic HBV to chronic HCV, with the latter showing levels comparable to confirmed AIH patients, suggests an immunological continuum representing different stages of liver immune activation.

The strong correlations between CXCL9 and CXCL13 observed exclusively in HCV and AIH groups indicate coordinated immune responses characteristic of autoimmune processes. These findings have significant clinical implications for differential diagnosis, patient monitoring, and therapeutic decision-making in chronic liver diseases.

Such results support the emerging concept of viral-induced autoimmune hepatitis and suggest that CXCL9 and CXCL13 may serve as valuable biomarkers for identifying patients at risk for autoimmune transition. This research contributes to understanding the complex interplay between viral infections and autoimmune phenomena, with potential implications for personalized treatment approaches in chronic liver diseases.

Future longitudinal studies incorporating these biomarkers could facilitate early identification of viral-to-autoimmune transition, potentially enabling preventive interventions and improving outcomes for patients with chronic hepatitis C infection.

## Acknowledgment

The authors express their heartfelt gratitude to the central laboratory personnel at the Gastrointestinal and Liver Disease Specialized Hospital in Al-Najaf, Iraq for their assistance during sample collection for the current investigation, as well as to all individuals who participated in this study.

